# Immunity to infections in arboviral vectors by integrated viral sequences: an evolutionary perspective

**DOI:** 10.1101/2020.04.02.022509

**Authors:** Cristina Crava, Finny S. Varghese, Elisa Pischedda, Rebecca Halbach, Umberto Palatini, Michele Marconcini, Annamaria Mattia, Seth Redmond, Yaw Afrane, Diego Ayala, Christophe Paupy, Rebeca Carballar-Lejarazu, Pascal Miesen, Ronald P. van Rij, Mariangela Bonizzoni

## Abstract

In the model organism *Drosophila melanogaster*, the PIWI-interacing RNA pathway contributes in silencing transposable elements (TEs) through smallRNAs (piRNAs), which arise from genomic loci (piRNA clusters) that contain sequences of previously-acquired TEs. As such, they are a functionally-immune archive of previous TE invasions that is passed to the offspring. In the arboviral vector *Aedes aegypti*, piRNA clusters contain TEs and endogenous viral elements from nonretroviral RNA viruses (nrEVEs) which produce piRNAs, supporting the hypothesis that nrEVEs are heritable immunity effectors. However, direct evidence that nrEVEs mediate adaptive immunity is lacking. Here, by using an analytic approach intersecting population genomics with molecular biology we demonstrate that the composition of piRNA clusters is modular through acquisition and absence of nrEVEs. We show that the genomes of wild-caught mosquitoes have a different set of nrEVEs than those annotated in the reference genome, including population-specific integrations. nrEVEs are not distributed in mosquito genomes only by genetic drift, but some show signs of positive selection. Moreover, by comparing natural mosquito populations expressing or lacking two newly characterised nrEVEs with high sequence complementarity to cell fusing agent virus, we show that nrEVEs confer antiviral immunity in ovaries against the cognate virus. Our results confirm that some nrEVEs have been co-opted for adaptive immunity to viral infections.

## INTRODUCTION

The PIWI-interacting RNA (piRNA) pathway, initially described as a silencing mechanism of transposable elements (TEs) in *Drosophila melanogaster* (Brennecke et al., 2007), shows striking functional similarities to the prokaryotic CRISPR-Cas system (Koonin and Makarova 2017). Both are sequence-based mechanisms that mediate immunity against foreign nucleic acids of which they form an “archive” in the genome (Marraffini and Sontheimer, 2010; Koonin and Makarova 2017; Kofler 2019). CRISPR RNAs (Clustered Regularly Interspaced Palindromic Repeats, crRNAs) are produced from short fragments of foreign nucleic acids previously integrated into the repetitive CRISPR locus of the host genome. crRNAs assemble with specific CRISPR-associated proteins (Cas) proteins to form complexes that bind and cleave any phage or plasmid bearing sequence complementarity to the crRNA (Amitai and Sorek et al., 2016).

piRNAs are a class of small RNAs of around 25-30 nt that guide PIWI proteins onto complementary target RNAs, resulting in gene silencing at the post-transcriptional or transcriptional level (Ozata et al., 2019). In *D. melanogaster*, piRNA precursors are generated from genomic loci, called piRNA clusters (Czech and Hannon, 2016). These regions are enriched for (remnants of) TE sequences and, as a consequence, cluster-derived piRNAs show sequence complementarity to TEs. It is thought that piRNA clusters act as traps for new TE invasions by horizontal transfer (Brennecke et al., 2007; Khurana et al., 2011; Parhad and Theurkauf, 2019). Thus, akin to CRISPR loci in prokaryotes for invading nucleic acids, piRNA clusters record the history of TE mobilization and provide a heritable source of piRNAs that silence active TEs present in the genome.

A single, large and uni-strand piRNA locus located in a heterochromatic region of the X chromosome, *flamenco,* controls TE movements in the somatic cells of the ovary in *D. melanogaster* (Koefler, 2019). *Flamenco* is composed of unique fragments from different TEs and changes in a single copy influence its regulatory properties (Zanni et al., 2013). TEs in the germline are repressed by the activity of several piRNA clusters, mostly dual-strand, distributed over multiple chromosomes and containing a different set of TE fragments than those in *flamenco* with instances of multiple fragments from the same TE, suggesting redundancy and tissue-specific regulation of certain TE classes (Malone et al., 2009; Duc et al., 2019).

Although the genome of the arboviral vector *Aedes aegypti* is richer in TE sequences than that of *D. melanogaster*, an early study suggested that its piRNA clusters are not enriched in TE fragments (Arensburger et al., 2011). We and others recently demonstrated that piRNA clusters of *Ae. aegypti* and *Ae. albopictus* are enriched in sequences from RNA viruses (Palatini et al., 2017; Whitfield et al., 2017; Palatini et al., 2020). These nonretroviral Endogenous Viral Elements (nrEVEs) or Nonretroviral Integrated RNA virus Sequences (NIRVS) derive primarily from insect-specific viruses (ISVs), which are non-reverse-transcribing RNA viruses that replicate exclusively in insects and are phylogenetically related to arboviruses (Roundy et al., 2017). ISVs are thought to be mainly maintained in mosquito populations by transovarial transmission (Agboli et al., 2019), perhaps explaining their predominance over arboviruses in insect genomes.

The physical contiguity between nrEVEs and TEs in piRNA clusters and the production of piRNAs from nrEVEs support the hypothesis that viral integrations contribute to mosquito immunity through piRNA-targeted processing of cognate viruses, and are adaptive (Palatini et al., 2017; Whitfield et al., 2017; Tassetto et al., 2019). Indeed, replication of recombinant Sindbis virus, engineered to carry a nrEVE sequence, was efficiently inhibited in *Ae. aegypti* Aag2 cells in a sequence and strand-specific manner (Tassetto et al., 2019), suggesting that nrEVEs have antiviral potential.

However, direct evidence that nrEVEs have antiviral activity and mediate *adaptive* immunity is lacking. Adaptation is the evolutionary process through which organisms adjust to a changing environment. In immunology, the term *“*adaptation” is used to indicate processes by which properties of immune cells or effectors are modified in response to environmental stimuli in a way that influence their subsequent responses to the same stimulus (Natoli and Ostuni, 2019). In vertebrates, the adaptive immune response generates memory of previous pathogens, resulting in more efficient targeting upon a second encounter, and depends on *de novo* generation of a diverse repertoire of antigen-specific receptors or effectors, which is highly dependent on the organism’s life history (Danilova, 2012; Bohem and Swann, 2014). Likewise, if nrEVEs are inherited adaptive immune effectors, it is expected that their distribution within mosquito genomes diversify according to previous pathogen encounters. Given the hundreds of nrEVEs of the *Ae. aegypti* genome, several of which are overlapping or corresponding to the same viral regions, it can also be hypothesized that nrEVEs represent ancient viral relicts, among which only few have been co-opted for antiviral functions (Katzourakis et al., 2017; Frank and Freschotte 2017).

Here, we applied computational and evolutionary approaches to whole-genome sequencing data of wild-caught *Ae. aegypti* mosquitoes to probe evolutionary and immunological adaptation of viral integrations. We identified novel nrEVEs in the genome of wild-collected mosquitoes mapping in and outside piRNA clusters and we showed that the landscape of nrEVEs is variable across populations. These results indicate that the composition of piRNA/nrEVE clusters can be modulated by environmentally-acquired viral fragments. Additionally, we show that not all nrEVEs evolve neutrally and signs of positive selection are clearly detected in a few viral integrations. Finally, we selected mosquito strains carrying newly identified nrEVEs and probed their effect on a subsequent infection with a cognate virus. We showed significantly reduced levels of virus from the secondary infection in ovaries, establishing antiviral immunity in a natural mosquito-virus system.

Overall these results establish that a number of nrEVEs of *Ae. aegypti* genomes have been maintained through selection and exert immunity functions. Additionally, we demonstrate that the landscape of viral integrations is modulable through new acquisition of viral integrations within and outside piRNA clusters.

## RESULTS and DISCUSSION

### Atlas of viral integrations in the *Aedes aegypti* genome

To obtain a genome-wide view of the complexity and distribution of nrEVEs in the highly contiguous AaegL5 reference genome, we used a previously-validated pipeline (Whitfield et al., 2017; Palatini et al., 2017). We annotated 213 nrEVEs, which derived from six viral families (Flaviviridae, Rhabdoviridae, Xinmoviridae, Phasmaviridae, Phenuiviridae and Mesoniviridae) plus several viruses that are still unclassified (fig. 1A, supplementary table 1). We also identified 39 Chuviridae-like nrEVEs which all derive from viral glycoprotein sequences and are completely embedded within long-terminal repeat (LTR) TEs, primarily of the Pao Bel family (supplementary table 2). LTR elements normally do not possess an envelope gene (*env*) but can acquire *env*-like genes from disparate viral sources and, through that, act similarly to infectious retroviruses (Hayward, 2017). For instance, *Drosophila* gypsy LTR elements gained infective properties after acquiring the *env* gene from baculoviruses (Malik et al., 2000). Given the peculiarity that all *Chuviridae*-like nrEVEs map within a subset of LTR TEs, this possibility cannot be excluded. It is beyond the scope of this work to test for infectious properties of Pao Bel elements carrying *Chuviridae*-like integrations and we excluded *Chuviridae*-like sequences from further analyses.

**Figure 1.**
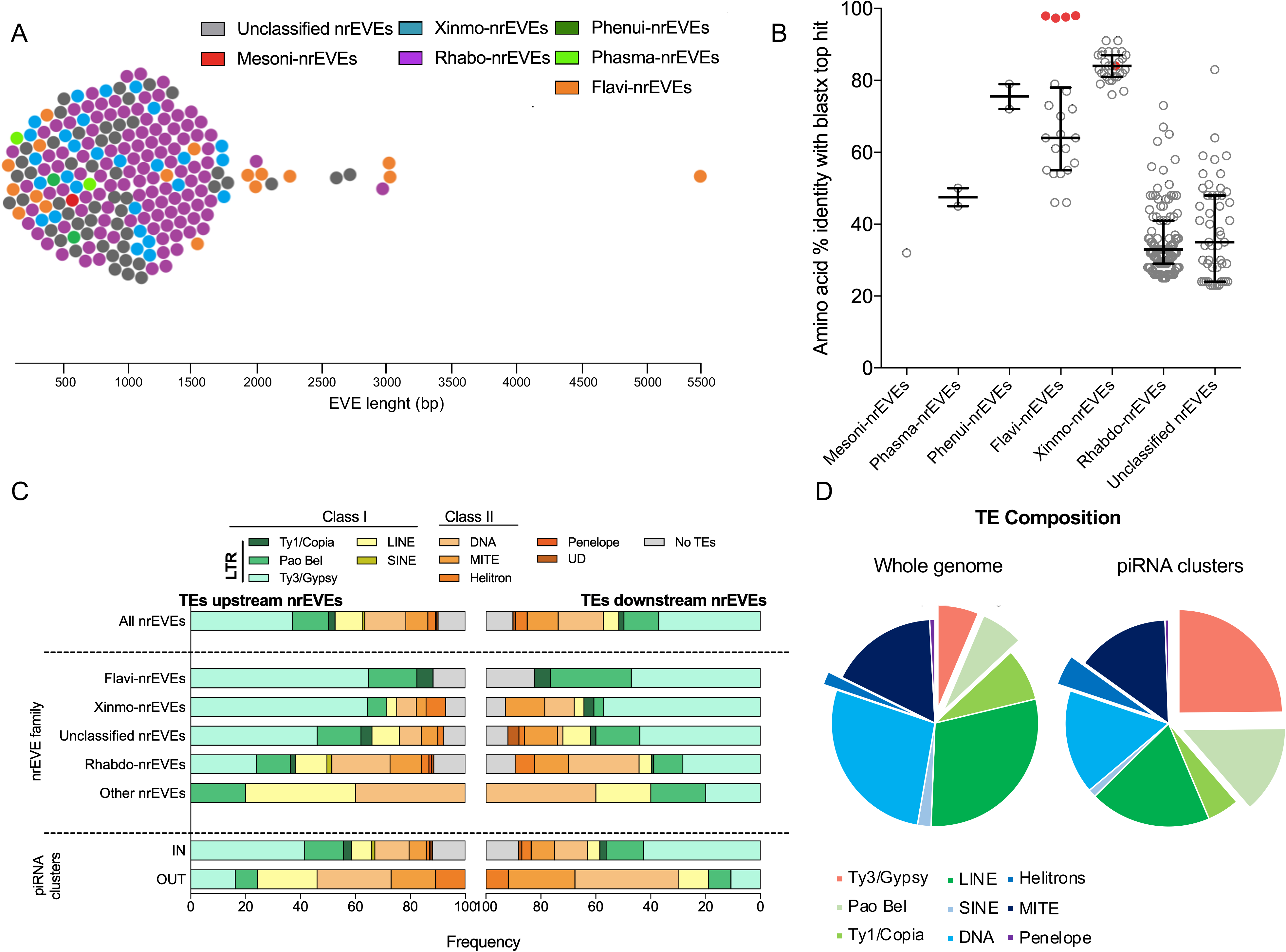
Atlas of nrEVE *Aedes aegypti*. **A)** Beeswarm plot showing nrEVEs identified in the *Ae. aegypti* genome (AagL5). Each dot represents a nrEVE, plotted according to its length and color-coded based on its viral origin. nrEVE lengths range from 136 to 5410 nt, with an average of 1028 nt. nrEVE lengths correlate with viral origin (Kruskal-Wallis test: H=15.87, 6 d.f., p=0.015), with sequences deriving from Flaviviridae (Flavi-nrEVE) being the longest. **B)** Scatter plot representing the amino acid identity of each nrEVE and its best hit retrieved by blastx searches against NR database grouped by viral family. Whiskers represent the median and the interquartile range. Red dots are the novel nrEVEs discovered in the geographical populations. **C)** Bar plots showing the type of the closest transposable element (TE) upstream and downstream of all nrEVEs (upper panel), nrEVEs grouped by their viral origin (middle panel), nrEVEs grouped by their location within (IN) or outside (OUT) piRNA clusters. Abbreviations: LTR (long terminal repeat), UD (unclassified TEs). **D)** Pie chart indicating the transposon composition of transposable elements (TEs) in the whole genome (left panel) or piRNA clusters (right panel).

Viral integrations distribute evenly on the three chromosomes, without enrichment at telomeric or centromeric regions and their identity with respect to the most similar virus ranges from 23 to 91% (fig. 1B; supplementary figure S1). Mapping against the reference viral genomes showed that nrEVEs derived from different portions of corresponding viral genomes (supplementary figure S2). nrEVEs from Rhabdoviridae (Rhabdo-EVEs) primarily originated from the nucleoprotein and glycoprotein sequences, whereas nrEVEs derived from Flaviviridae (Flavi-EVEs) arise mainly from regions of the viral genome encoding for nonstructural proteins, primarily NS1 and NS2 (supplementary table 1).

### The piRNA profile of viral integrations

Given the high number of nrEVEs in the *Ae. aegypti* genome, we asked whether the presence of multiple sequences corresponding to the same region of the viral genome would favor amplification of the piRNA signal. To answer this question, we first annotated piRNA clusters in the *Ae. aegypti* AaegL5 genome assembly and then mapped deep-sequencing data from somatic and germline tissues (ovaries) to viral integrations. We identified 1158 clusters occupying around 1% of the *Ae. aegypti* genome (supplementary table 3). From these, 108 and 38 clusters exclusively expressed in the germline or somatic tissues, respectively (fig. 2A). 176 nrEVEs were located within piRNA clusters spanning 219 kb (1.6% of the total piRNA cluster length of 14Mb) (fig. 2B). Five piRNA clusters harbor >10 nrEVEs (fig. 2C), mostly from different viral families, including all Flavi-EVEs located in piRNA clusters (i.e. piRNA clusters 2q44.4.) and are active in both soma and ovaries where they tend to produce piRNAs from one dominant strand (uni-strand clusters, fig. 2D, supplementary table 3). Dual-strand piRNA clusters are mostly active in the germline (fig. 2E, supplementary figure S3). As previously observed, nrEVEs are tightly associated with LTR retrotransposons (fig. 1C), especially Ty3/gypsy and Pao Bel retrotransposons (Palatini et al., 2017; Whitefield et al., 2017). The association between Ty3/Gypsy and nrEVEs does not depend on the distance between the two elements, but it is linked to the location of a viral integration within or outside piRNA clusters (supplementary tables 1,3). Almost half (47.4%) of TEs associated with nrEVEs within piRNA clusters are Ty3/Gypsy retroelements whereas they represent only 13.9% of the TEs associated with nrEVEs that map outside piRNA clusters (Fisher’s exact test, p<0.001). LTR retrotransposons, especially Ty3/gypsy and Pao Bel retrotransposons, are enriched in piRNA clusters compared to the genome (fig. 1D). Thus, the association between nrEVEs and Ty3/Gypsy elements is biased by the overall enrichment of Ty3/Gypsy elements in piRNA clusters (Fisher exact test p<0.001).

**Figure 2.**
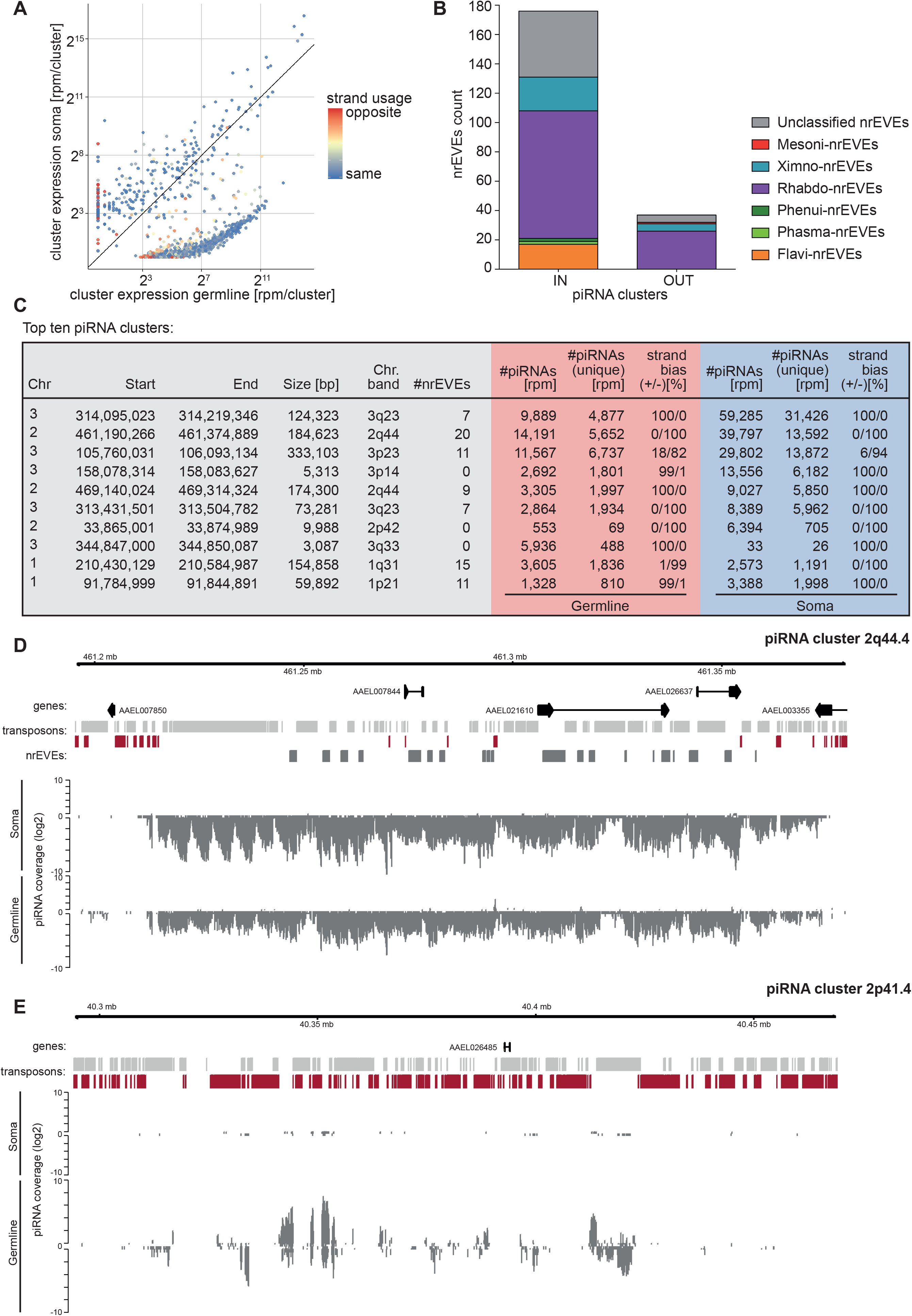
*Aedes aegypti* piRNA clusters. **A)** Expression of piRNA clusters in germline and somatic tissues. piRNA coverage per million mapped small RNAs (rpm) plus a pseudo-count of 1 is plotted in order to include values of zero. Color indicates the likelihood of a cluster being expressed with the same strand bias in both tissues. **B)** Bar plots showing the distribution of nrEVEs from different viral families within (IN) and outside (OUT) piRNA clusters. **C)** Table listing the top-ten most highly expressed piRNA clusters with at least 5% uniquely mapping piRNA reads. **D)** Coverage plot of a piRNA cluster with strand bias towards expression from one strand (uni-strand). **E)** Coverage plot of a piRNA cluster without strong strand bias (dual-strand). Log2 coverage in both germline (ovaries) and somatic tissues is shown. Genes are indicated with black arrows, transposons are indicated with light gray (plus strand) or red (minus strand) boxes, and nrEVEs are depicted with dark gray boxes (plus strand).

piRNAs are found mapping to all nrEVEs, with the exception of Meso1, Xin23, Xin24, Rha17, Rha45, Rha60 and Rha69, consistent with their location outside piRNA clusters (supplementary table 1). Within each viral family, there are both nrEVEs that share 100% nucleotide identity (hereafter called overlapping nrEVEs) as well as nrEVEs that correspond to the same viral region with nucleotide identity below 60%, likely representing integrations of sequences of different viruses. In contrast, overlapping nrEVEs may represent independent integrations of the same virus or, more likely, duplication of the nrEVE sequence after integration into the mosquito genome. For overlapping nrEVEs, we cannot unambiguously map piRNAs to a single viral integration (Supplementary Table 4). nrEVE-derived piRNAs are not evenly distributed, but spike in distinct portions across the integration independently of the number of corresponding viral integrations (fig. 3). For Flavi-EVEs, piRNA hotspots are in regions corresponding to NS1, NS2 and NS3 sequences of Flaviviruses (fig. 3A). Despite Flavi-EVEs being distributed across five piRNA clusters that are active both in the soma and the germline, their piRNA profile is more pronounced in the soma (supplementary table 3; fig. 3A). In contrast, piRNA signals for Rhabdo-EVEs were mostly detected in the germline, and spanned the nucleoprotein (N), glycoprotein (G) and polymerase (L) sequences (fig. 3B). Discontinuous piRNA profiling within a piRNA cluster was already noticed in *Drosophila* as a result of both the local and long-range sequence environment (Muerdter et al., 2012).

**Figure 3.**
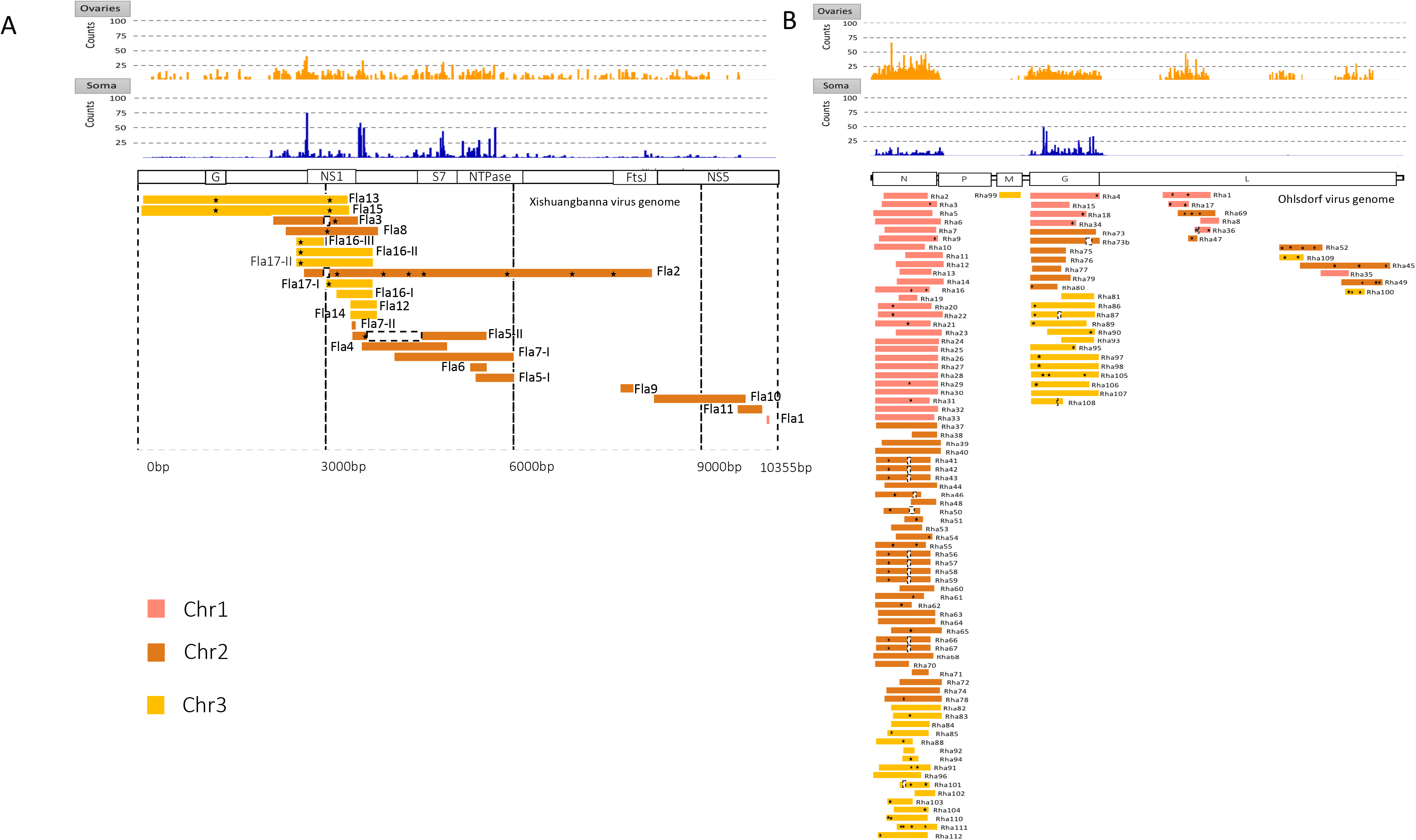
Distribution and piRNA coverage of nrEVEs on reference viral genomes. **A)** Flaviviridae-derived nrEVEs (Flavi-nrEVE) aligned to the Xishuangbanna flavivirus genome (NC_034017.1) **B)** Rhabdoviridae-derived nrEVE (Rhabdo-nrEVE) aligned to the Ohlsdorf virus genome (KY768856). Rha73, Flavi5, Flavi7, Flavi16, Flavi17 are composed of repeated and not contiguous parts of the viral genome and thus have been fragmented to overlap to the corresponding viral location. Stars indicate stop codons or small INDELs that interrupt the viral open reading frame and dotted white boxes indicate large deletions that generate stop codons. Top panels indicate piRNAs mapping to the indicated positions in soma (orange) and ovaries (blue), respectively.

Overall, these results provide an overview of the nrEVE landscape in *Ae. aegypti* and their potential for piRNA-mediated antiviral activity. The biased piRNA profile along viral integrations suggests that *some* nrEVEs may have been selected for antiviral functions.

### Wild mosquitoes harbor novel viral integrations

We next analyzed nrEVE distribution in the genomes of sixteen individual mosquitoes from five geographic populations each, searching specifically for novel nrEVEs that are absent from the *Ae. aegypti* reference genome. The samples included three populations from Africa, one from Mexico, and one from American Samoa. Throughout Africa, *Ae. aegypti* occurs predominantly as a darker form, called *Ae. aegypti formosus* (Aaf), which feeds on animals and uses natural water collections including tree holes for the larval development. Outside of Africa, a lighter, domesticated form of *Ae. aegypti*, called *Ae. aegypti aegypti* (Aaa), occurs, which feeds on humans and uses anthropogenic containers as larval breeding sites (Crawford et al., 2017). The divergence between Aaf and Aaa is estimated to have occurred between 5,000-10,000 years ago prior to the global Aaa expansion (Crawford et al., 2017; Rose et al., 2020).

A total of five novel nrEVEs were identified in our geographic samples (fig. 4, supplementary dataset 1). Four have similarity to the insect-specific flavivirus cell fusing agent virus (CFAV), with a nucleotide identity of over 97% with the Galveston reference strain (fig. 1B). The fifth novel nrEVE has similarity to another insect specific virus, *Aedes* anphevirus (AeAV) with 85% identity. CFAV-derived integrations correspond to different genomic regions of the viral genome and three of them likely arose from recombination of CFAV sequences which are not contiguous in the CFAV genome (fig. 4A). The longest novel viral integration (i.e. CFAV-EVE-1) is a 3909 bp sequence composed by three regions of the CFAV genome, with one encompassing the complete viral CDS corresponding to the NS1 protein. A second novel CFAV-like integration (CFAV-EVE-2) is 1259 bp long and is composed of two parts of similar length that include parts of the E protein and the glycoprotein NS1, and the central part of NS3, respectively. A third novel CFAV-like integration (CFAV-EVE-3) is 734 bp-long and is composed of four parts (two of them inserted in reverse orientation compared to CFAV genome) spanning part of NS2A and NS2B, NS4A, NS4B and NS5 coding sequences. The shortest CFAV-like integration (CFAV-EVE-4) is a 328 bp-long sequence, corresponding to part of the NS2A coding region of CFAV (fig. 4A). The novel AeAV-like integration contained a single genomic sequence that corresponds to a portion of the viral nucleoprotein (fig. 4A). All novel CFAV-like integrations had no polymorphism nor indels among all geographic mosquitoes that have the respective integration. PCR amplified products from each novel nrEVE were obtained from additional mosquitoes and sequenced, providing a molecular validation of their bioinformatic-based identification (supplementary figure S4).

**Figure 4.**
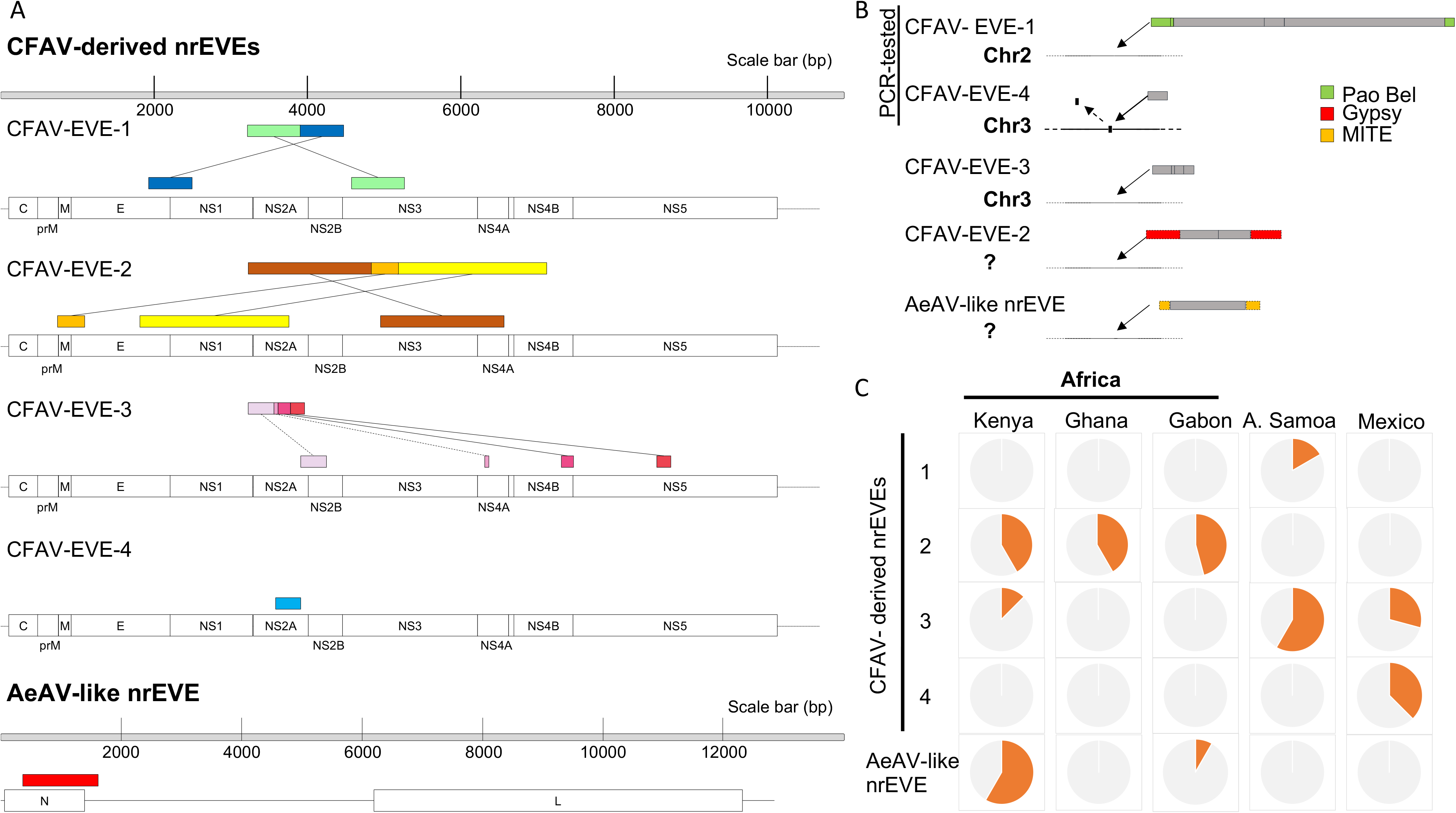
Novel viral integrations in wild collected *Aedes aegypti* mosquitoes. **A)** Scheme of the novel viral integrations with similarity to Cell fusing agent virus (CFAV) and Aedes anphevirus (AeAV) identified in the genome of wild-collected mosquitoes. CFAV-EVEs are mapped to the genome of CFAV Galveston strain (NCBI Reference Sequence: NC_001564.2) and AeAV-like CFAV to AeAV strain MRL-12 genome (MH037149.1). Dotted lines represent part of CFAV-EVE-3 that integrated in the opposite direction compared to the CFAV genome. **B)** Scheme of the integration points and flanking sequences of the novel nrEVEs; nrEVE sequences are represented by grey boxes. **C)** Frequency distribution of novel viral integrations tested with PCR in 24 mosquitoes from each site (Kenya, Ghana, Gabon, American Samoa and Mexico).

For three of the five novel nrEVEs, chromosomal integration sites could be deduced by *de novo* assembly of sequence reads, which were further confirmed by PCR and Sanger sequencing for two of them (fig. 4B). CFAV-EVE-1 inserted in chromosome 2, at position 294,058,716, together with three fragments of a Pao Bel LTR-TE (Pao Bel 277 in Matthews et al., 2018) giving a hybrid sequence longer than 5900 bp. CFAV-EVE-3 inserted in chromosome 3, at position 137,741,235. CFAV-EVE-4 inserted between genomic positions 106,043,272 and 106,043,299 on chromosome 3, leading to loss of the intervening 27-nt sequence (Fig. 4B). CFAV-EVE-4 is located in piRNA cluster 3p23.4, which is highly active in both soma and germline and hosts other viral integrations, including eight Rhabo-EVEs, one Xinmo-EVE, and two unclassified nrEVEs. Besides acquisition of CFAV-EVE-4, we also observed the absence of the Xin18, Rha89 and Rha88 nrEVEs from piRNA cluster 3p.23.4 in the genomes of African mosquitoes (Fig.5). These results are the first proof that the composition of nrEVEs within piRNA clusters can be naturally modulated in wild mosquitoes.

**Figure 5.**
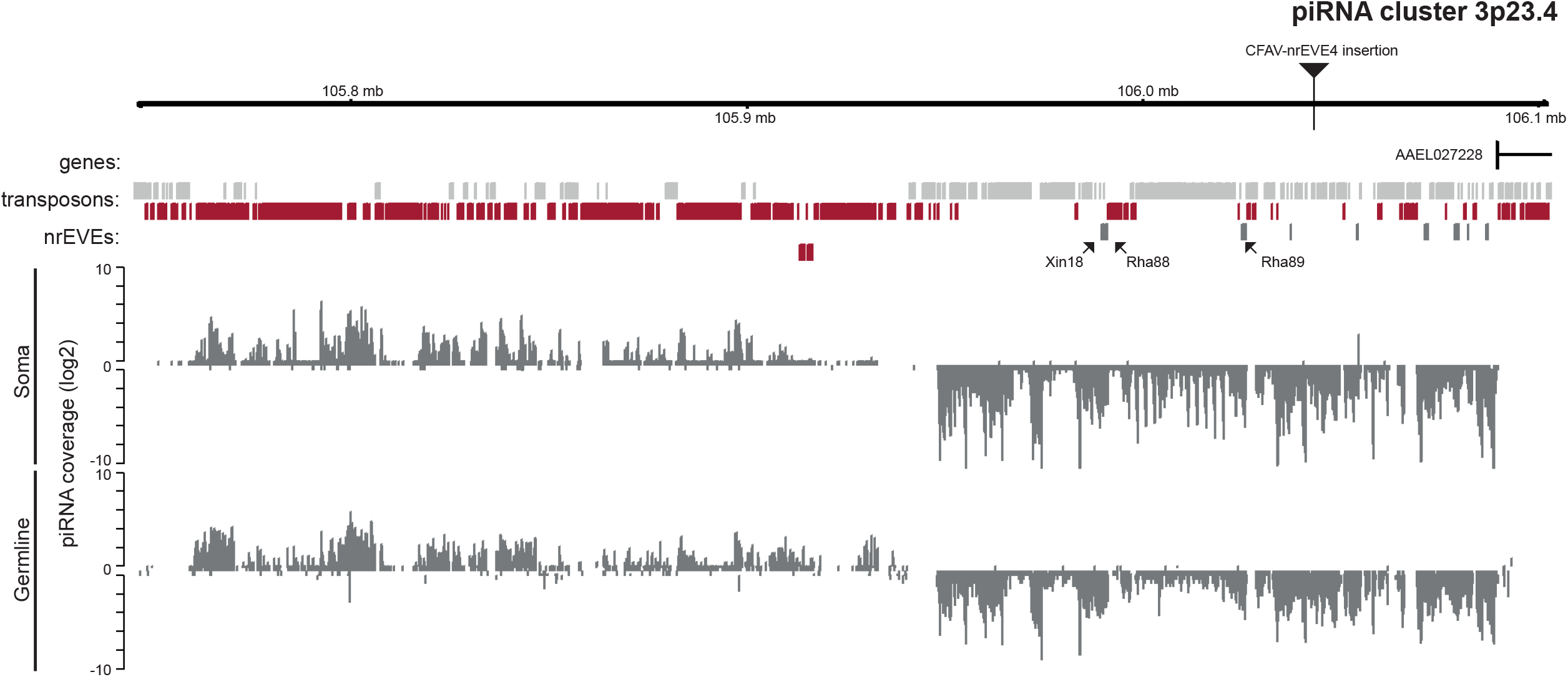
piRNA clusters can be modulated by acquisition of viral sequences. Coverage plot of piRNA cluster 3p23.4. Genes are indicated with black arrows, transposons and nrEVEs are indicated with gray (plus strand) or red (minus strand) boxes. Arrows indicate variably-distributed nrEVEs which are present only in some individuals.

Regions flanking CFAV-EVE-2 correspond to Gypsy245, a LTR Gypsy TE that is present in multiple copies in the *Ae. aegypti* genome (Matthews et al., 2018). The repetitive nature of the flanking regions of CFAV-EVE-2 along with the lack of knowledge on how variable the TE landscape is across *Ae. aegypti* populations prevent the characterization of the CFAV-EVE-2 integration site. However, a clean signal was detected in linked-read (10X) data in two mosquitoes of a presumably single-copy viral insertion around 461MB on chromosome 2, at the far end of the Q arm. Unexpectedly, in a third mosquito, the CFAV-EVE-2 sequence was integrated in chromosome 3 (supplementary figure S4). The AeAV-like EVE was flanked by mTA element 38c, a DNA transposon of the miniature inverted–repeat transposable elements (MITEs) family that is present in multiple copies in the genome, thus preventing the precise mapping of this novel AeAV-like integration.

### Novel viral integrations are population-specific

Novel viral integrations were not fixed in any population and displayed a population-specific pattern. CFAV-EVE-2 was detected only in samples from Africa, CFAV-EVE-1 only in America Samoa, and CFAV-EVE-4 only in Mexico at frequencies of 43.1%, 16.7% and 37.5%, respectively (fig. 4C). CFAV-EVE-3 and AeAV-like EVE were present in multiple populations at frequency varying from 8.3% to 58.3%. As the Kenyan and Gabon samples are the ancestral Aaf form, these results indicate that CFAV-EVE-3 is an ancestral integration that occurred before the split between Aaf and Aag. We cannot exclude that the absence of CFAV-EVE-2 in the new world samples here analysed is due to drift. First records of *Ae. aegypti* in Mexico are from the seventeenth century and *Ae. aegypti* populations from Oceania and surrounding islands are derived from American populations, probably through navigation routes well-established by the nineteenth century (Powell et al., 2018), suggesting CFAV-EVE-1 and CFAV-EVE-4 are local integrations.

### Wild mosquitoes have a variable landscape of viral integrations

To gain insights into nrEVE evolution, we analyzed patterns of occurrence of viral integrations in the geographic samples and their polymorphism in relation to slow- and fast-evolving *Ae. aegypti* genes (Pischedda et al., 2019). We reasoned that, if viral integrations result from fortuitous events and are viral fossils, their distribution should be governed by drift and their polymorphism should evolve at a neutral rate (Aswad and Katzourakis, 2012; Katzourakis, 2013). If viral integrations behave like TE fragments within piRNA clusters and are antiviral, their overall genomic landscape is expected to be variable across host genomes depending on pathogen exposure and nrEVE sequences may co-evolve with those of pathogens (Zanni et al. 2013; Goriaux et al. 2014; Asif-Laidin et al. 2017; Frank and Freschotte 2017).

A total of 70 viral integrations appear to be shared across all populations, representing a conserved core of old viral integrations (fixed EVE) (fig. 6A). There is a strong bias for Flavi-EVEs among the fixed integrations, with 12 out of 13 tested Flavi-EVEs (92%) being detected in all individuals and in all populations (supplementary table 1). The frequency patterns of variably distributed (VD)-EVEs separates samples according to their geographic origin (fig. 6B). However, population frequencies of VD-EVEs are not statistically-different based on the likelihood ratio test (LRT) (supplementary table S5).

**Figure 6.**
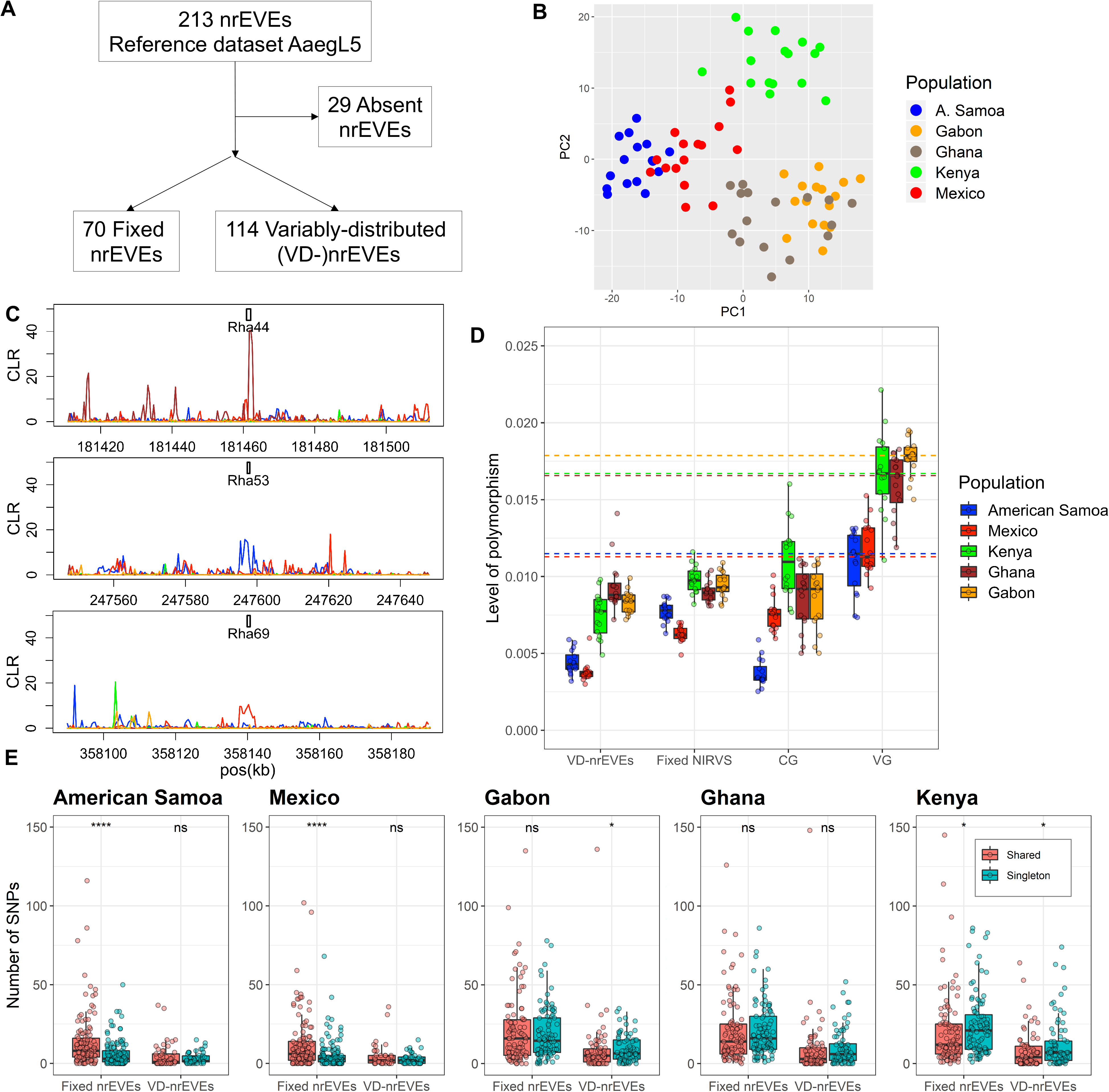
Geographic variation in nrEVE distribution. **A)** Outline of the distribution of nrEVEs observed in all populations (fixed) and nrEVEs observed only in some populations (variably distributed, VD-nrEVEs). **B)** Convex logistic principal component analysis (PCA) of VD-nrEVE frequencies based on their geographic origin. Each dot indicates an individual mosquito, color-coded based on geographic location. **C)** Composite likelihood ratio (CLR) signal around the indicated nrEVEs in mosquito populations color-coded as in panel B. Increased CLR signal around the nrEVE is indicative of positive selection. **D**) Whisker plots comparing the level of nucleotide polymorphism among *Ae. aegypti* populations in conserved genes (CG), fast-evolving genes (VG), fixed and VD-nrEVEs. Each dot represents the average value of an individual mosquito, boxes span the interquartile range, marked lines within the boxes represent the median and whiskers represent the minimum and the maximum. Dotted lines are color-coded according the population they represent and depict the median value of the level of polymorphism of fast-evolving genes. **E)** Comparison of the number of singletons (*i.e.* SNPs found only in one individual) versus SNPs in VD- or fixed-nrEVE in each population. Statistical differences were established by the Wilcoxon rank sum test ((ns not significant, * *p*-value < 0.05, ** *p*-value < 0.01, *** *p*-value < 0.0001)).

### Sign of positive selection on viral integrations

The spectrum of polymorphism within, and flanking, a DNA sequence can be used to infer its evolutionary history under the premises of the theory of neutral molecular evolution (Kimura, 1983). The theory states that most of the polymorphism at the DNA-sequence level is a neutral balance between two forces: mutation, which introduces new variants, and genetic drift, which randomly eliminates polymorphism. A number of statistical tests are available to compare the observed polymorphism of a region to that expected under neutral evolution given assumptions of population size, demography, random mating and recombination (for a review, see Booker et al., 2017; Pavlidis and Alachiotis, 2017). Deviations from expectations under the null hypothesis of neutral evolution are interpreted as signs of selection. An increase in the frequency of a variant, associated with reduced variability in the neighboring region due to genetic hitchhiking, generates a hard-selective sweep, which indicates positive or adaptive selection. The power of detecting signatures of hard selective sweeps using single nucleotide polymorphisms (SNPs) is sufficiently strong when they occurred less than ~0.1 Ne generations ago (Kim and Stephan 2000; Przeworski 2002), where Ne is the effective population size. Given that *Ae. aegypti* has around 11 generations per year and an Ne of roughly 500 (Saarman et al. 2012), this translates in the ability to detect only very recent hard sweeps, in the order of ~4 to 8 years. Thus, selective sweeps detected in sequence data are likely to be population specific and, in the case of beneficial nrEVEs associated with locally circulating viruses. Signatures of hard sweeps were detected for Rha44 in Ghana, Rha69 in Mexico and Rha53 in American Samoa (fig. 6C). Rha44 and Rha69 are fixed in the populations where hard sweep was predicted; Rha53 is close to fixation in American Samoa (Freq= 0.938), supporting the hypothesis that the increase in frequency of these three nrEVEs is likely due to positive selection.

Adaptive evolution at viral integrations was also tested by estimating Tajima’s D statistics genome-wide and verifying if nrEVEs map in windows with significantly negative Tajima’s D values (Koefler et al., 2012; Rech et al., 2019). Tajima’s D compares the polymorphism and the segregating sites in a region under the premises of the neutral evolution theory: a low Tajima’s D value occurs in presence of an excess of low frequency variants resulting from a bottleneck or a selective sweep; on the contrary a value of Tajima’s D significantly higher than 0 indicates a scarcity of rare variants, interpreted as balancing selection or recent population admixture (Stephan, 2019). Consistent with results of hard sweep, Rha69 showed significantly negative Tajima’s D values in Mexico (Fig. 6). Rha44 and Rha53 showed lower than chromosome-average Tajima’s D values in Ghana and American Samoa, albeit values were not statistically significant. Among these three nrEVEs, only Rha44 maps in a piRNA cluster, 2p21.18, which is active only in the germline (supplementary table 3). Additional sixteen nrEVEs showed significantly negative Tajima’s D values in different populations, with Rha2, Rha35, Rha86 and Un33 consistently in two African populations (fig. 7). These Rhabdo-EVEs derive from different viral regions and different viruses and all map in piRNA clusters active in both somatic and germline tissues (supplementary Tables 1, 3).

**Figure 7.**
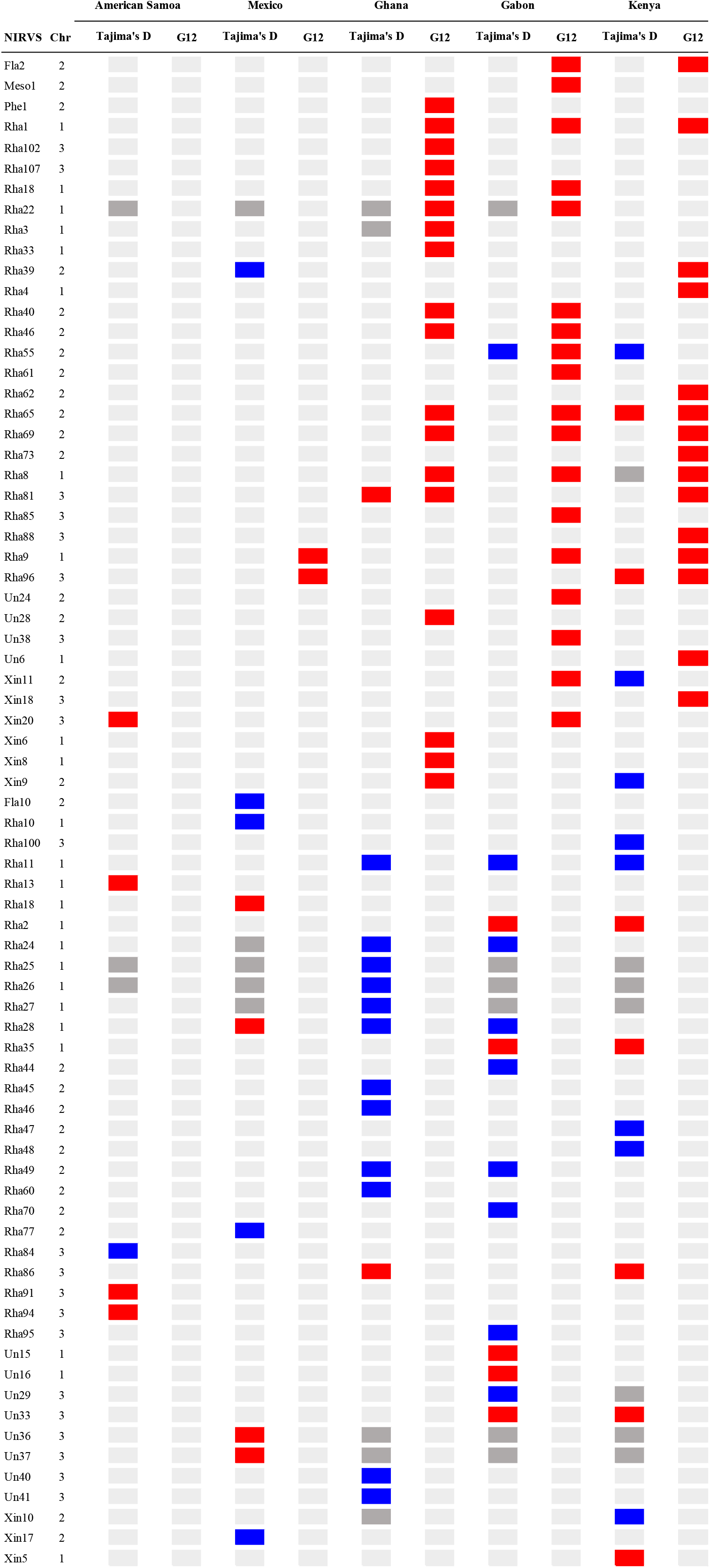
Fixed-nrEVE with signal of selection. Tajima’s D and G12 values in the indicated nrEVEs in geographic populations of *Ae. aegypti*. Red indicates higher Tajima’s D or lower G12 values than the cutoffs for each chromosome in each population (Supplementary Table 9). Blue boxes indicate Tajima’s D values higher than the 5% cutoffs for each chromosome in each population. Dark and light gray boxes indicate instances of non-significant tests or tests that could not be calculated, respectively.

An allele may be present in a population along with other variants and segregate neutrally, until an environmental change arises that favors its segregation. This situation will result in a soft selective sweep, detectable through *ad hoc* statistics such as the G12 and G1/2 method (Harris et al., 2018; Garud et al., 2015). We calculated the G12 and G1/2 statistics genome-wide and verified whether nrEVEs occurred in windows with the top 15% most extreme G12 values, following Rech et al., 2019. Signatures of soft sweep were identified in 36 nrEVEs, primarily in African populations; among these viral integrations, Rha65 and Rha96 in Kenya and Rha81 in Ghana also had significantly low Tajima’s D values. We hypothesized that this result is driven by the presence of different haplotypes, of which one at low frequency. In Ghana, the signature of soft sweep at Rha55 was accompanied by a significantly high Tajima’s D value (fig. 6), supporting that Rha55 is under positive balancing selection in this population. These viral integrations are found both outside piRNA clusters (*i.e.* Rha55 and Rha81) or within germline piRNA clusters (*i.e.* Rha65 in 2q22.16 and Rha96 in 3p14.2). To further confirm that some nrEVEs are adaptive, we compared their polymorphism with that of two previously validated sets of fast-evolving and conserved *Ae. aegypti* genes, respectively (Pischedda et al., 2019). If nrEVEs are biological inactive relics and have reached fixation by genetic drift, they should accumulate mutations at the host mutation rate. Consistently, viral integrations that are variably distributed in tested populations are expected to be younger, thus less polymorphic than fixed ones. Contrary to this expectation, we found that VD-nrEVEs showed similar polymorphisms as fixed ones, especially in the oldest African populations (fig. 6D). Additionally, polymorphism and Tajima’s D values were lower in viral integrations than in fast-evolving genes (supplementary figure S5). These results are against expectation of nrEVEs slowly drifting to high frequency and evolving neutrally.

We also analyzed the distribution of SNPs *versus* singletons (*i.e*. SNPs found only in one mosquito) across viral integrations in each population. An excess of singletons is indicative of negative selection (Bourgeois and Boissinot, 2019). No significant excess of singletons was observed in any population (fig. 6E); in contrast, Rhabdo- and Flavi-EVEs showed statistically more SNPs than singletons, primarily in Mexico and American Samoa (supplementary figure S5). This result holds for viral integrations both within and outside piRNA clusters. Overall, our analyses of nrEVE polymorphism showed that not all nrEVE evolve similarly and few bear clear signs of positive selection.

### Novel nrEVE and viral infection

*Aedes aegypti* mosquito lines containing the novel integrations CFAV-EVE2 and CFAV-EVE3 were generated from field-caught CFAV EVE-positive mosquitoes from Kenya. In parallel, an outcrossed control line lacking any of the novel CFAV EVEs was established. To test our hypothesis that CFAV EVEs can provide immunity to incoming cognate viruses, we tested whether the CFAV-EVE-2 and CFAV-EVE-3 mosquito populations are less susceptible to CFAV infection as compared to the control population. As insect specific flaviviruses are thought to be vertically transmitted (Agboli et al., 2019), we separately analyzed female germline tissue (ovaries) and carcasses.

To this end and to facilitate dissection of the mosquito ovary, we infected mosquitoes in the gonadotropic cycle, during which ovaries develop and mature towards oviposition. At 16 h post-feeding, blood-fed mosquitoes were intra-thoracically injected with 500 TCID_50_ of CFAV. Based on CFAV growth kinetics performed in blood-fed *Ae. aegypti* mosquitoes (supplementary figure S6), we selected day 2 post-infection to dissect ovaries. CFAV viral RNA loads were analyzed from the carcasses and ovaries of each genotype-confirmed ‘CFAV-EVE2’, ‘CFAV-EVE3’ and ‘Control’ mosquito. No significant difference was observed in viral RNA levels between the carcasses of the three different groups. However, CFAV viral RNA loads were significantly higher (more than 10-fold) in the ovaries of the control group as compared to those from the ‘CFAV-EVE2’ (P < 0.01), and ‘CFAV-EVE3’ (P < 0.001) groups (fig. 8). Thus, CFAV-EVE containing mosquitoes are more resistant to infection with a CFAV strain bearing high sequence similarity to the EVE sequence. These data provide evidence for a small RNA-based adaptive immunity that exists in nature against a naturally circulating virus.

**Figure 8.**
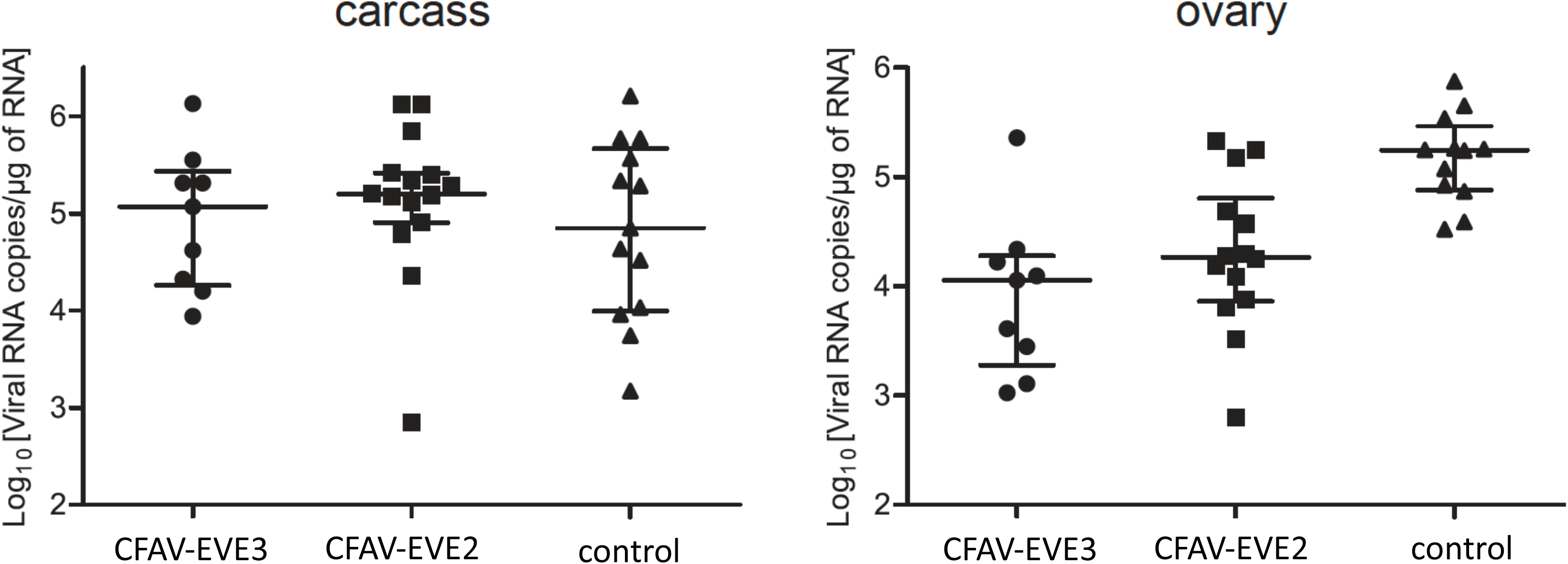
CFAV replication is restricted in novel CFAV EVE-containing mosquitoes. Blood-fed *Ae. aegypti* Kenyan strain mosquitoes with the indicated novel CFAV nrEVEs were intra-thoracically injected with 500 TCID_50_ of CFAV (strain Rio Piedras 02). Viral load was determined in carcasses (left panel) and ovaries (right panel) dissected at 2 days post infection by RT-qPCR. Viral loads were estimated based on a standard curve of a plasmid containing the CFAV NS3 gene segment. Each symbol represents a single mosquito, with median and interquartile range indicated. Differences in viral load were tested on log-transformed data using ANOVA, followed by a Dunnett’s post hoc test. *** P < 0.001; ** P < 0.01.

## CONCLUSIONS

Endogenous viral elements have been hypothesized to mediate antiviral immunity in mosquitoes, yet direct evidence for this hypothesis is lacking. Starting from a thorough annotation of the organization and distribution of nrEVEs and piRNA clusters in the improved reference genome of *Ae. aegypti* (AaegL5) we here demonstrated that wild-caught mosquitoes have a variable landscape of nrEVEs and provide evidence that some nrEVEs are under positive selection. Moreover, we provide direct evidence that nrEVEs mediate antiviral immunity in ovaries. Our combined population genomics and experimental approach thus provides strong evidence for an antiviral function of nrEVEs. Our genome annotation of both nrEVEs and piRNA clusters represent an additional valuable resource for the vector biology community and pinpoint differences among nrEVEs.

Among the novel viral integrations identified, CFAV-EVE-4, is unique to mosquitoes from Mexico and maps in one of the most active *Ae. aegypti* piRNA cluster, 3p23.4. This cluster also hosts additional viral integrations that show signs of soft sweep (i.e. Rha86, Un33 and Rha91). To our knowledge this is the first demonstration that the composition of a mosquito piRNA cluster is modulated through the natural acquisition of a viral sequence. We observed additional piRNA clusters with a different composition of nrEVEs across geographic populations. For instance, the unistrand piRNA cluster 2q44.4, which is active both in the soma and germline, contains the nrEVEs Fla6, Rha76 and Xin10, which occur at a frequency < 0.5 in American Samoa, Ghana and Mexico, respectively. Interestingly, 2q44.4 contains additional viral sequences, a portion of which have 100% sequence identity with Fla6, Rha76 and Xin10 (i.e. Fla2, Fla5 and Fla7, Rha75 and Rha77 and Xin11), suggesting rearrangements among viral integrations contribute in defining the composition of piRNA cluster 2q44.4.

We were able to derive strains from wild-collected eggs that selectively bear two of the newly identified viral integrations CFAV-EVE-2 and CFAV-EVE-3. These viral integrations provide protection from subsequent infection with cognate viruses in ovaries, but not in carcasses. We propose that these nrEVEs produce piRNAs that poise the piRNA machinery to target incoming viruses with high sequence complementarity. The reason why antiviral activity is only observed in ovaries is currently unclear, but may be due to differential expression of PIWI proteins or associated proteins (Akbari et al., 2013).

A thorough analysis of nrEVE polymorphism clearly showed that not all viral integrations evolve neutrally. This result and the absence of orthologs between nrEVEs of *Ae. aegypti* and those of *Ae. albopictus* prevents dating integration events (Aiewsakun and Katzourakis, 2015). We can, however, distinguish between ancestral integrations occurring in both our *Ae. aegypti formosus* (i.e. Ghana, Kenya, and Gabon samples) and *Ae. aegypti aegypti* samples (Mexico, American Samoa) vs integrations detected exclusively in new world *Ae. aegypti* mosquitoes, which are thus younger than 10,000 years. The first group includes CFAV-EVE-3 in which no polymorphism was observed across our geographic samples and for which we established antiviral activity in ovaries. The second group includes Rha63 and CFAV-EVE-4 or Rha101 and CFAV-EVE-1, exclusively detected in American Samoa and Mexico mosquitoes, respectively. These data demonstrate that acquisition of viral sequences is a continuous, albeit rare event.

Signs of positive selection were primarily identified in Rhabdo-EVEs and in a population-specific manner. Rhabdoviruses are a family of viruses with a broad host range including plants, insects, fish, reptiles, mammals and crustaceans (Dietzgen et al., 2017). Recent metagenomics projects identified multiple insect-specific viruses from different phylogenetic taxa of the family Rhabdoviridae in mosquitoes (Ohlund et al., 2019; Atoni et al., 2019) and pervasive endogenization of rhabdoviruses into the genomes of arthropods and plants (Katzourakis and Gifford, 2010; Chiba et al., 2011; Fort et al., 2012). These results point to a long-standing association between rhabdoviruses and mosquitoes that merits further investigation.

Overall, our data clearly demonstrate that nrEVEs are a complex component of the mosquito repeatome being maintained through both drift and selection and that some nrEVEs have been co-opted for antiviral immunity. The distribution of nrEVEs across mosquito genomes, their representation of different portions of corresponding viral genomes associated with hot spots in their piRNA profiles suggest that nrEVEs are organised through a redundant system and their antiviral function may not be a universal feature, but depend on sequence identity, the genomic context in which they occur, and the portion of the viral genome from which they derive.

## MATERIALS AND METHODS

### nrEVE annotation in the *Ae. aegypti* genome assembly AaegL5

The latest *Ae. aegypti* genome assembly (AaegL5) (Matthews et al., 2018) was used to identify nrEVEs using iterative standalone blastx searches. A viral database was composed at the end of August 2018 using ssRNA, dsRNA and unclassified viral amino acid sequences available in the NCBI refseq viral database with host flag invertebrates plus two *Aedes anphevirus* sequences available from Genbank (ID AWW13507 and AWW13504) (supplementary table S6). Initial blastx was run using the viral database and the *A. aegypti* genome as query with a conservative e-value of 10^−6^.

Overlapping nrEVEs were then merged using the EVE FINDER pipeline (Whitfield et al., 2017). A second blastx search (e-value 10^−6^) with newly identified viral integrations as a query was run against the whole refseq amino acid database. Predicted viral integrations that had the highest identity to eukaryotic genes were manually discarded. Viral taxonomy was then assigned to each nrEVE based on the top hit retrieved by blastx searches against non-redundant (NR) database, which was used to extract the corresponding viral family from the NCBI taxonomy database. When a viral family could not be extracted from the NCBI database, that viral integration was annotated as “unclassified”. The BED file containing all transposable elements (TEs) annotated in the *Ae. aegypti* genome (AaegL5 assembly) (Matthews et al., 2018) was parsed with BEDTools to find TEs overlapping each viral integration (Quinlan and Hall, 2010). Maximum observed overlap between nrEVEs and TEs was 100bp and we removed the overlapping region from the annotation of the nrEVE sequence. BEDTools was further used to identify the closest genomic feature (i.e. a TE, another nrEVE or a coding sequence) to each nrEVE. The presence of Chuviridae-like EVEs within a TE was analyzed using a combination of BEDtools closest and BEDtools intersect. A custom script was used to assess the completeness of the TEs, relative to the TE landscape of *Ae. aegypti* (Matthews et al., 2018). Potentially coding ORFs were identified using GeneMark with a Heuristic Approach (Besemer and Borodovsky, 1999).

### piRNA cluster annotation in the *Ae. aegypti* genome assembly *AaegL5*

Clusters were annotated on the current *Ae. aegypti* genome assembly (AaegL5) (Matthews et al., 2018) using two publicly available small RNAseq datasets from blood fed female *Ae. aegypti* germline (ovaries) and female somatic tissues (carcasses) (SRR5961506 and SRR5961505, respectively) (Lewis et al., 2018). Adapters were clipped from reads with cutadapt (v1.18) (Martin, 2011), and the reads were then mapped with bowtie without allowing mismatches (v1.2.2) (Langmead et al., 2009). Thereby, ambiguously mapping reads were randomly distributed across all possible mapping positions (--best --strata -M 1 –seed 123) or discarded to retain only uniquely mapping reads associated with single-copy piRNA *loci* (-m 1).

piRNAs clusters were annotated analogously to the approach used in *D. melanogaster* (Brennecke et al., 2007). Briefly, only the first 5’ nucleotide of each piRNA sized read (23-32nt) was used and normalized to the total number of mapped piRNAs per million (ppm) within each library. The genome was scanned with non-overlapping 5 kb sliding windows, applying and optimizing various threshold values such as piRNA density per window, unambiguity of piRNA mapping, as well as size and minimal piRNA density per cluster. For the final cluster annotation all windows with 10 or more ppm (supplementary figure S2) and a maximum distance of 5 kb were merged into a single cluster. Clusters were required to contain at least 5 single-copy (unique) piRNA loci, and to be covered by at least 5 uniquely mapping piRNAs per million (supplementary figure S2). The 5’ and 3’ end of the cluster were defined by the most distant piRNAs within the merged windows. Finally, all clusters that were either very small (< 1kb) or had a very low read coverage (average coverage < 10 ppm/kb) were filtered out (supplementary figure S2). Initial piRNA cluster annotation was performed separately for ovary and somatic tissues, after which the results were merged to reach the final cluster annotation.

### Analysis of piRNA production from nrEVE

The same small RNAseq datasets (SRR5961506 and SRR5961505) used for piRNA cluster prediction were mapped to the *Ae. aegypti* genome (AaegL5 assembly) using bowtie with a minimum seed match of 18 nt. Aligned reads were filtered by length using BBMap reformat.sh (https://sourceforge.net/projects/bbmap/), keeping only piRNA sized reads (23-32nt) (Czech and Hannon, 2016). BEDTools Intersect (Quinlan and Hall, 2010) removed reads mapping outside annotated nrEVEs. Finally, all reads with 100% identity were collapsed with fastx-toolkit (http://hannonlab.cshl.edu/fastx_toolkit/) and used for quantification. Due to sequence similarity and overlap among nrEVEs, it is impossible to quantify reads mapping uniquely to single nrEVEs. To avoid any bias, we used custom scripts to quantify each piRNA sized sequence in each original fastq file and then identified all the viral integrations to which each piRNA could be mapped. Counts for each experiment were normalized based on the library size by Quantile-to-Quantile Normalization as implemented in edgeR (Robinson et al., 2009).

### Geographical samples

*Aedes aegypti* mosquitoes were sampled as adults by BG-sentinel traps or as larvae from tires and backhoe buckets in the summer and fall of 2017 in Tapachula (Mexico), Franceville (Gabon), Larabanga (Ghana), M’barakani village near Rabai (Kenya), and Tafuna Village, Tutuila Island (America Samoa). Larvae were reared to adulthood *in situ* and ethanol-preserved adults were shipped to the University of Pavia (Italy). Additionally, *Ae. aegypti* eggs were collected using ovitraps in M’barakani (Kenya) and Franceville (Gabon) in December 2018 and were hatched at the University of Pavia (Italy). At adulthood, mosquitoes were checked for *Flavivirus* infection using degenerate primers (Crochu et al., 2004), but no infection was detected. Mosquitoes were reared under constant conditions, at 28°C and 70-80% relative humidity with a 12h/12h light/dark cycle. Larvae were reared in plastic containers, at a controlled density to avoid competition for food. Fish food (Tetra Goldfish Gold Colour) was provided daily. Adults were kept in 30 cm^3^ cages and fed with cotton soaked in 0.2 g/ml sucrose as a carbohydrate source. Adult females are fed with defibrinated mutton blood (Biolife Italiana) using a Hemotek blood feeding apparatus. Based on the dark coloring of the abdomen, all *Ae. aegypti* samples from Africa appeared to be *Ae. aegypti formosus* (Mattingly, 1957).

### Genome sequence generation

Genomic DNA was extracted individually from 16 adult mosquitoes from each population with the Promega Wizard Genomic DNA Purification Kit, according to the manufacturer’s protocol. Individual DNA libraries were prepared with TruSeq DNA PCR-free reagents and sequenced to a minimum 20x coverage (average 24x) on the Illumina HiSeq X Ten platform by Macrogen to generate paired-end 150 nt reads. Raw reads were trimmed with Trimmomatic v0.38 (Bolger et al., 2014). Whole genome sequencing data have been submitted at NCBI SRA under BioProject PRJNA609256.

### Bioinformatic pipeline to detect reference nrEVEs

The presence or absence of each viral integration characterized from the *Ae. aegypti* reference genome (AaegL5 assembly) was analyzed in genomic resequencing of individual mosquitoes using an in-house bioinformatic pipeline, which also allows for sequence polymorphism analysis (Pischedda et al., 2019). The pipeline allows to detect the presence of nrEVEs in each tested individual, but not to distinguish between heterozygote and homozygote status. Hence, we approximated allele frequency as the number of viral integrations normalized by the total number of individuals. Because of the stringency used in the call for nrEVE presence (i.e. minimum five reads with at least 30 consecutive nucleotides with mapping quality higher than 20) (Pischedda et al., 2019), a short sequence that shares sequence identity with a longer one could be erroneously called absent in all individuals tested because of reads shared with the longer nrEVE. For this reason, a total of 29 viral integrations, which were called absent in all individuals, were excluded from further analyses (supplementary table 1).

### Identification of novel nrEVEs via linked-read sequencing

We investigated a set of 28 sequences, representing a broad geographic and genetic distribution of *Ae. aegypti*, that had been sequenced with linked-read (10X) sequencing that generated libraries in which reads that derived from a single strand of DNA are tagged with a unique barcode (Redmond et al. 2020). Comparison of common barcodes allows inference of proximity between reads up to 80kb apart. To identify novel nrEVE integration sites, each readset was aligned to novel nrEVEs to identify any reads that might derive these viral integrations; following detection of novel nrEVEs, reads that were linked to these sequences were then aligned to the AaegL5 genome identifying the sequence flanking the viral integration. Background signal can derive from misalignment or multiple occupancy of 10X droplets; for each 1MB window we calculated the sequence covered by flanking reads and positions of viral integrations were determined as those within the 0.999^th^ percentile (supplementary figure S4).

### Identification of novel nrEVEs

We used Vy-PER followed by custom scripts to search for novel viral integrations (Forster et al., 2015). Because Vy-PER uses BLAT (Kent, 2002) which recognises sequences of 95% and greater sequence similarity of at least 40 bp in length, we restricted our search for novel viral integrations to 167 viral species already identified as part of mosquito virome (Supplementary Table 7). Viral integration candidates were checked for the absence of eukaryotic sequences. Additionally, all reads whose viral region had a dinucleotide percentage higher than 80% and a viral length shorter than 50 bp were removed. Trinity was used for *de novo* reassembly of the putative viral integration (Grabherr et al., 2011). Finally, these *de novo* assembled viral integrations and their flanking regions were re-mapped to the *Ae. aegypti* genome using blastn to characterize the exact point of integration.

### Molecular analysis to confirm novel nrEVEs

Novel nrEVEs and their flanking regions were amplified by PCR and Sanger sequenced to confirm their presence in the mosquito genome. PCR was carried out with the DreamTaq Green PCR Master Mix (ThermoFisher) using 1 μl of 1:10 dilution of the DNA that had been used for next-generation sequencing. Amplified bands were purified with ExoSAP-IT kit (ThermoFisher) and Sanger sequenced (Macrogen, Madrid, Spain). Sequences were analyzed with Bioedit (Hall, 1999). After confirming the identity of each viral integration, we used PCR to analyze the distribution of these nrEVEs in the tested populations. Primers used are listed in Supplementary Table 8.

### Analysis of nrEVE polymorphism

nrEVE polymorphism was analyzed at two levels. First, the frequency distribution of each nrEVE was analyzed by investigating its presence or absence in each tested mosquito. Second, sequence polymorphism of each nrEVE was estimated in single individuals. nrEVE distribution across geographical samples was visualized using convex logistic PCA from the R package logisticPCA (Landgraf and Lee, 2015). Heterogeneity in the occurrence of nrEVEs among populations was evaluated using a maximum likelihood procedure adapted from a study on the distribution of TEs in *D. melanogaster* (Gonzalez et al., 2008), assuming that sampled populations are panmictic. Thus, the data for each nrEVE can be described as {*m*_1_, *m*_2_} where *m1* is the number of individuals in which a nrEVE was present, independently from its genotypic status, and *m2* is the number of individuals in which that nrEVE is absent. The log-likelihood of observing such data conditional to the frequency *p* is:

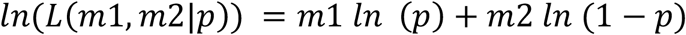

The *L*(*p*) is maximized at the value 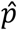:

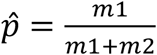

To determine whether the frequencies were different among populations an LRT test was used that compares two models. Under the null hypothesis, we assumed that the frequencies of viral integrations were the same in all populations and estimated 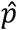 using combined data from all populations. Under H1, we assumed that nrEVE frequencies were different among populations and estimated 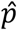 for each population, separately. We then calculated maximum log-likelihood for both 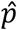 and they were compared as:

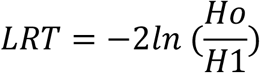

Heterogeneity was detected when LTR was greater than 9.49 corresponding to 5% of the *χ*^6^ test with four degrees of freedom.

Sequence polymorphism of each nrEVE was estimated by analyzing their SNPs and calculating the level of polymorphism (LoP), as previously described (Pischedda et al., 2019).

### Signature of selection

We applied different methods to test for signatures of positive selection. The presence of hard selective sweep was predicted using SweeD (Pavlidis et al., 2013) on a window size of 100 Kb (50 Kb upstream and 50 Kb downstream of the nrEVE) using SNP and INDEL dataset called with SAMtools/BCFtools (Li, 2011). For each data set, the composite-likelihood ratio (CLR) was calculated over a grid of 250, which resulted in estimates over ∼400 bp. CLR estimates were visualized with R (Team, 2014). Candidate nrEVEs harboring a signature of a selective sweep were selected when their CLR values were higher than the 99^th^ percentile of their corresponding window distribution. Signatures of soft sweep were predicted using the G12 statistics implemented in the SelectionHapStat software (Garud et al., 2015) after having identified SNP variants using Freebayes (Garrison and Marth, 2012). The H12_2H1.py script was run using 50 SNPs as the window size for each of the three *Ae. aegypti* chromosomes in each population, with overlaps of 25 bp for each window. We used small and overlapping window sizes to avoid biases from recombination; linkage disequilibrium in *Ae. aegypti* is estimated between 52-67 Kb (Matthews et al., 2018). Windows in the top 15% most extreme G12 values were selected and analyzed for the presence of viral integrations (Rech et al., 2019).

Selection in fixed nrEVEs was also tested by calculating Tajima’s D values (Tajima, 1989), a procedure previously applied to test for selection in fixed TEs (Kofler et al., 2012; Rech et al., 2019). Tajima’s D values were calculated in non-overlapping 500 bp windows using vcftools v.0.1.15 (Danecek et al., 2011). The approach of Rech et al., (2019) was used to identify windows with significantly low Tajima’s D values. Average Tajima’s D values were first calculated per chromosome and population and windows with Tajima’s D values lower than the 5th percentile of the whole chromosome distribution were then analyzed (Supplementary Table 9). Finally, significant windows were screened for the presence of nrEVEs.

### Viruses

CFAV strain UVE/CFAV/2002/PR/Rio Piedras 02 (Ref-SKU: 0001v-EVA68) isolate was obtained from the European Virus Archive. Virus stocks were prepared and titrated on *Ae. albopictus* C6/36 cells (Cook et al., 2009; Moureau et al., 2007).

### Cells

*Ae. albopictus* C6/36 cells were cultured in L15 medium (Gibco) supplemented with 10 % heat-inactivated FBS (Gibco), 2 % tryptose phosphate broth (Sigma), 50 U/ml penicillin (Gibco) and 50 μg/ml streptomycin (Gibco).

### CFAV replication kinetics

*Ae. aegypti* Liverpool strain mosquitoes (BEI Resources, NR-48921) were starved for 24 h and subsequently fed with human blood (Sanquin Blood Supply Foundation) using a Hemotek membrane feeding system. The mosquitoes were injected at 16 h post-blood-feeding with 500 TCID_50_ of CFAV Galveston strain in full L15 medium using a Nanoject-II Auto-Nanoliter Injector (Drummond) with a 3.5-inch glass capillary needle (Drummond). Ovaries were dissected from the surviving CFAV-infected mosquitoes on days 0, 3 and 5 post-infection. Ovaries and the carcasses from each individual mosquito were processed as described below.

### Mosquito blood-feeding and infection

Female adult mosquitoes from ‘CFAV-EVE2, ‘CFAV-EVE3’ and ‘Control’ groups, maintained on 10 % sucrose solution, were starved for 24 h before feeding them with human blood (Sanquin Blood Supply Foundation) using a Hemotek membrane feeding system. Engorged females were then sorted into separate containers and provided with sucrose solution again. 16 h post-blood-feeding, mosquitoes were injected with 500 TCID_50_ of CFAV Galveston strain in full L15 medium using a Nanoject-II Auto-Nanoliter Injector (Drummond) with a 3.5-inch glass capillary needle (Drummond). As controls, blood-fed mosquitoes from each group were mock-infected with L15 medium alone. At 2 days post-infection, ovaries were dissected from surviving CFAV-infected and mock-infected mosquitoes from each group. Dissected ovaries and the remaining carcasses were homogenized using 1 mm diameter zirconium beads (BioSpec) in RNA-Solv reagent (Omega Bio-Tek) and total RNA was isolated from these samples according to the manufacturer’s instructions.

### Genotyping

Due to the heterozygotic nature of our CFAV EVE-containing ‘CFAV-EVE2’ and ‘CFAV-EVE3’ populations, a pair of legs from each mosquito was kept aside for EVE genotyping. Two legs from each dissected mosquito were homogenized using 1 mm zirconium beads (BioSpec) in nuclease-free water. Homogenate was used as a template for PCR using internal primers for ‘CFAV-EVE2’ and ‘CFAV-EVE3 sequences (Supplementary Table 8). All mosquito samples were cross-tested for both integrations.

### Viral load

RNA isolated from mosquito ovaries and carcasses were subjected to DNase treatment (Ambion) and used to synthesize cDNA with random hexamers using the TaqMan Reverse Transcription reagent (Applied Biosystems). Quantitative real-time PCR was performed using GoTaq qPCR (Promega) BRYT Green Dye-based detection and CFAV NS3-specific primers. The following primers were used. CFAV NS3 Fw 5’–TTATGGACCGGGATGACATT–3’ and CFAV NS3 Rev 5’–GCGGTCATCAACGTATTGTG–3’. Primers do not have sequence complementarity to the sequence of both integrations. CFAV NS3 PCR product was cloned into a pGEM-3Z vector using the *Bam*HI restriction site and used as quantitative copy number standard for viral load estimation. The time course experiment (Supplementary Figure 6) was analyzed using CFAV NS1 Fw 5’– GCAGCGGCGCTTTTGTGTGG–3’ CFAV NS1 Rev 5’–GCACTGCAAGGCATCCTCAC–3’. Differences in viral load were tested on log-transformed data using ANOVA, followed by a Dunnett’s post hoc test (Sigmaplot) in GraphPad Prism.

## Acknowledgements

We thank Mark Schmaedick, from the American Samoa Community College, for providing access to mosquitoes from American Samoa. We thank Francesca Scolari and Lino Ometto from the University of Pavia for mosquito maintenance and fruitful discussions. CFAV, strain Rio Piedras 02, was kindly provided by Dr. Remi Charrel, Aix Marseille University, through the European Virus Archive (EVA) funded by the European Union, 7^th^ Framework programme. *Ae. aegypti* Liverpool strain mosquitoes were kindly provided by the NIH/NIAID Filariasis Research Reagent Resource Center, distributed by BEI Resources, NIAID, NIH.

## SUPPLEMENTARY MATERIAL

**Supplementary Figure S1. Chromosomal distribution of nrEVEs of *Aedes aegypti*.** Distribution of viral integrations across the three *Ae. aegypti* chromosomes: each arrow may refer to a single nrEVE or several nrEVEs located in close proximity. Total number of viral integrations is shown under each chromosome. Number of viral integrations for each family is shown between parentheses. **A**) nrEVEs reference dataset; **B**) *Chuviridae*-like nrEVEs

**Supplementary Figure S2. Distribution of nrEVEs on reference viral genomes.** Xinmoviridae-derived EVEs (Xinmo-EVE) were aligned on Aedes anphevirus genome (NCBI accession MH037149); Phenuiviridae-derived EVEs (Phenui-EVEs) to Phasi Charoen-like virus segment S (NC_038263); Phasmaviridae-derived EVEs (Phasma-EVEs) to Culex phasma-like virus segment M (MF176243) and Mesoniviridae-derived EVEs (Mesoni-EVEs) to Nse virus ORF 1a (NC_020901). Stars indicate stop codons or small INDELs that interrupt the viral ORF, dotted white boxes indicate large deletions that generate stop codons.

**Supplementary Figure S3. piRNA cluster annotation. A)** Percentage of piRNAs residing within clusters (left panel), or fraction of the genome that is occupied by piRNA clusters (right panel) that can be annotated when using different threshold values for a minimal density of piRNAs per 5 kb window. **B)** Percentage of piRNAs within clusters when using different thresholds for single-copy piRNA loci a cluster must contain. These clusters were annotated with a minimal piRNA density of 10 piRNAs per 5kb and a maximum distance of 5 kb between windows. **C)** piRNA density and cluster size for clusters with 5 single-copy piRNA loci and at least 5 unique piRNA reads per million. Clusters that fall below 1 kb and/ or 10 ppm/kb are removed in later steps. Colors indicate the strand-usage of the individual clusters (percentage of piRNAs derived from the dominant strand).

**Supplementary Figure S4. Molecular and bioinformatics analysis of novel viral integrations. A)** Results of PCR amplifications of five novel viral integrations identified in single mosquitoes from five geographic populations of *Ae. aegypti* (pres: presence; abs: absence). Primers for CFAV-EVE-4 were designed in the regions flanking the integration resulting in the identification of heterozygotes individuals. **B)** Detection of CFAV-EVE-2 insertion site in *Ae. aegypti* genomes that had been sequenced with linked-read (10X). Linked-reads (10X) were aligned to CFAV-EVE-2 to identify nrEVE-derived reads; flanking reads were identified via 10X barcode and aligned to the AaegL5 assembly. The proportion of sequence covered by flanking reads within 1 mb windows was compared and windows containing CFAV-EVE-2 integration determined as those above the 0.999^th^ percentile (dashed line). Three samples are here shown to have the CFAV-EVE-2 integration, two at the distal end of Chr 2Q and one at the proximal end of Chr 3Q. The distribution of these sites in two different populations, and the lack of uniform integrations within each population may indicate selective maintenance of CFAV-EVE-2.

**Supplementary Figure S5. Population genomics and evolution of nrEVEs. A)** Geographic distribution of nrEVEs. The number of variably distributed (VD) NIRVS is plotted with respect to their frequency in each population. In American Samoa, most VD-NIRVS were present at high frequency (>50%), indicating their distribution is driven by drift and a reduced efficiency of negative selection. **B)** Venn-diagram showing the number of viral integrations shared across the tested populations. **C)** Departures from neutrality as measured by the Tajima’s *D* test statistic. Values were estimated for each of the 50,000 bp long genomic windows (n = 223,455 to 228,456, depending on the population), 132-134 of which contained conserved genes (CG), 159-164 contained fast evolving genes (VG) and 189-197 contained nrEVEs. **D)** Distribution of singletons (*i.e.* SNPs found only within one individual) versus SNPs in Flavi- and Rhabdo-EVEs across the five tested populations. Differences between Tajima’s D averages, as well as between the number of singletons and SNPs, were tested by the Wilcoxon rank sum test (ns not significant, * *p*-value < 0.05, ** *p*-value < 0.01, *** *p*-value < 0.0001).

**Supplementary Figure S6 CFAV growth kinetics in *Aedes aegypti* mosquitoes.** Blood-fed *Ae. aegypti* Liverpool strain mosquitoes were intra-thoracically injected with 500 TCID_50_ of CFAV (strain Rio Piedras 02). Viral RNA load was determined by RT-qPCR from ovaries and carcasses, dissected at the indicated days post-infection.

**Supplementary Table 1**. Atlas of nrEVEs in the *Aedes aegypti* genome (AaegL5 assembly).

**Supplementary Table 2**. *Chuviridae*-like sequences integrated into the *Aedes aegypti* genome.

**Supplementary Table 3**. List of *Aedes aegypti* piRNA clusters (AaegL5 assembly).

**Supplementary Table 4**. List of nrEVEs-piRNAs

**Supplementary Table 5**. nrEVEs with skewed geographic distribution: A) nrEVE with the highest LRT indicating differences in the frequency patterns across populations; B) population-specific viral integrations; C) viral integrations absent in one of the tested populations, but present in others.

**Supplementary Table 6**. List of viral species (and proteins) used for the characterization of nrEVEs in the *Aedes aegypti* genome (AegL5 assembly)

**Supplementary Table 7.** Viral database used to identify novel viral integrations.

**Supplementary Table 8.** List of PCR primers used to detect presence/absence of novel nrEVEs.

**Supplementary Table 9.** Cut-off values for Tajima’s D and G12 statistics.

**Supplementary dataset 1**. Fasta sequences of novel nrEVEs.

**Supplementary Table 1**. Atlas of nrEVEs in the *Aedes aegypti* genome (AaegL5 assembly).

**Supplementary Table 2**. *Chuviridae*-like sequences integrated into the *Aedes aegypti* genome.

**Supplementary Table 3**. List of *Aedes aegypti* piRNA clusters (AaegL5 assembly).

**Supplementary Table 4**. List of nrEVEs-piRNAs

**Supplementary Table 5**. nrEVEs with skewed geographic distribution: A) nrEVE with the highest LRT indicating differences in the frequency patterns across populations; B) population-specific viral integrations; C) viral integrations absent in one of the tested populations, but present in others.

**Supplementary Table 6**. List of viral species (and proteins) used for the characterization of nrEVEs in the *Aedes aegypti* genome (AegL5 assembly)

**Supplementary Table 7.** Viral database used to identify novel viral integrations.

**Supplementary Table 8.** List of PCR primers used to detect presence/absence of novel nrEVEs.

**Supplementary Table 9.** Cut-off values for Tajima’s D and G12 statistics.

**Supplementary dataset 1**. Fasta sequences of novel nrEVEs

